# C1 CAGE detects transcription start sites and enhancer activity at single-cell resolution

**DOI:** 10.1101/330845

**Authors:** Tsukasa Kouno, Jonathan Moody, Andrew Kwon, Youtaro Shibayama, Sachi Kato, Yi Huang, Michael Böttcher, Efthymios Motakis, Mickaёl Mendez, Jessica Severin, Joachim Luginbühl, Imad Abugessaisa, Akira Hasegawa, Satoshi Takizawa, Takahiro Arakawa, Masaaki Furuno, Naveen Ramalingam, Jay West, Harukazu Suzuki, Takeya Kasukawa, Timo Lassmann, Chung-Chau Hon, Erik Arner, Piero Carninci, Charles Plessy, Jay W Shin

## Abstract

Single-cell transcriptomic profiling is a powerful tool to explore cellular heterogeneity. However, most of these methods focus on the 3’-end of polyadenylated transcripts and provide only a partial view of the transcriptome. We introduce C1 CAGE, a method for the detection of transcript 5’-ends with an original sample multiplexing strategy in the C1™ microfluidic system. We first quantified the performance of C1 CAGE and found it as accurate and sensitive as other methods in C1 system. We then used it to profile promoter and enhancer activities in the cellular response to TGF-β of lung cancer cells and discovered subpopulations of cells differing in their response. We also describe enhancer RNA dynamics revealing transcriptional bursts in subsets of cells with transcripts arising from either strand within a single-cell in a mutually exclusive manner, which was validated using single molecule fluorescence in-situ hybridization.

## Introduction

Single-cell transcriptomic profiling can be used to uncover the dynamics of cellular states and gene regulatory networks within a cell population(Trapnell, 2015; Wagner, Regev and Yosef, 2016). Most available single-cell methods capture the 3’-end of transcripts and are unable to identify where transcription initiates. Instead, capturing the 5’-end of transcripts allows the identification of transcription start sites (TSS) and thus the inference of the activities of their regulatory elements. Cap analysis gene expression (CAGE), which captures the 5’-end of transcripts, is a powerful tool to identify TSS at single nucleotide resolution(Shiraki *et al*., 2003; Carninci *et al*., 2006). Using this technique, the FANTOM consortium has built an atlas of TSS across major human cell-types and tissues(Forrest *et al*., 2014), analysis of which has led to the identification of promoters as well as enhancers in the human genome(Andersson *et al*., 2014; Hon *et al*., 2017). Enhancers have been implicated in a variety of biological processes(Lam *et al*., 2014; Li, Notani and Rosenfeld, 2016), including the initial activation of responses to stimuli(Arner *et al*., 2015) and chromatin remodeling for transcriptional activation(Mousavi *et al*., 2013). In addition, over 60% of the fine-mapped causal noncoding variants in autoimmune disease lay within immune-cell enhancers (Farh *et al*., 2015), suggesting the relevance of enhancers in pathogenesis of complex diseases. Enhancers have been identified by the presence of balanced bidirectional transcription producing enhancer RNAs (eRNAs), which are generally short, unstable and non-polyadenylated (non-polyA)(Andersson *et al*., 2014). Single molecule fluorescence *in situ* hybridization (smFISH) studies have suggested that eRNAs are induced with similar kinetics to their target mRNAs but that co-expression at individual alleles was infrequent(Rahman *et al*., 2016). However, the majority of enhancer studies have been conducted using bulk populations of cells meaning that the dynamics of how multiple enhancers combine to influence gene expression remains unknown.

The majority of single-cell transcriptomic profiling methods(Picelli, 2017) rely on oligo-dT priming during reverse transcription, which does not capture non-polyA RNAs transcripts (e.g. eRNAs). The recently developed RamDA-seq(Hayashi *et al*., 2018) method uses random priming to capture the full-length non-polyA transcripts including eRNAs. However, this method is not strand-specific and unable to pinpoint transcript 5’-ends; thus, it cannot detect the bidirectionality of eRNA transcription and cannot confidently distinguish reads derived from the primary transcripts of their host gene (i.e. intronic eRNAs). Methods are typically implemented for a specific single-cell handling platform (e.g. microwell, microfluidics or droplet-based platforms)(Picelli, 2017), because each platform imposes strong design constraints on the critical steps of cell lysis and nucleic acid handling. The proprietary C1™ Single-Cell Auto Prep System (Fluidigm) uses disposable integrated fluidic circuits (IFCs) and provides a registry of publicly available single-cell transcriptomics methods (Supplementary Table 1), which can be customized. Previously, we introduced nanoCAGE(Plessy *et al*., 2010), a method requiring only nanograms of total RNA as start material, based on a template switch mechanism combined with random priming to capture the 5’-ends of transcripts independent of polyA tails in a strand-specific manner. Here we develop C1 CAGE, a modified version of nanoCAGE customized to the C1 system to capture the 5’-ends of transcripts at single-cell resolution.

Current single-cell methods are usually limited in the number of samples that can be multiplexed within the same run. Thus, experimental designs requiring multiple replicates and different conditions are prone to batch effects, confounding biological information with the technical variation of each experiment(Tung *et al*., 2017). To mitigate batch effects, we took advantage of the transparency of the C1 system to encode multiple cells perturbation states in a single run by fluorescent labeling and imaging.

We apply this method to investigate the response to TGF-β in A549 cells, an adenocarcinomic human alveolar basal epithelial cell line. TGF-β signaling plays a key role in embryonic development, cancer progression, host tumor interactions and driving epithelial-to-mesenchymal transition (EMT)(Massagué, 2008; Ikushima and Miyazono, 2010). We examine the response to TGF-β in A549 cells to uncover dynamically regulated promoters and enhancers at single-cell resolution. We observed an asynchronous cellular response to TGF-β in sub-populations of cells. We also investigated the dynamics of enhancer transcription at single-cell resolution with validation by smFISH. Our results suggest transcriptional bursting of enhancers as reflected by high expression of eRNAs in a few cells. Also, while in pooled cells enhancers show bidirectional transcription, within single-cells transcription at enhancers is generally unidirectional—i.e. transcription on the two strands seems to be mutually exclusive.

## Results

### Development of C1 CAGE

We developed the C1 CAGE method, based on nanoCAGE(Plessy *et al*., 2010), C1 STRT Seq(Islam *et al*., 2014) and C1 RNA-seq(Wu *et al*., 2014), implementing reverse transcription with random hexamers followed by template switching and pre-amplification (Figure 1a). The cDNA is tagmented and the 5’-end of cDNA is specifically amplified by index PCR. The resulting library is sequenced from both ends, with the forward reads identifying the 5’-end of the transcript at single nucleotide resolution and the reverse read identifying downstream regions of the matching transcript.

**Figure 1:**
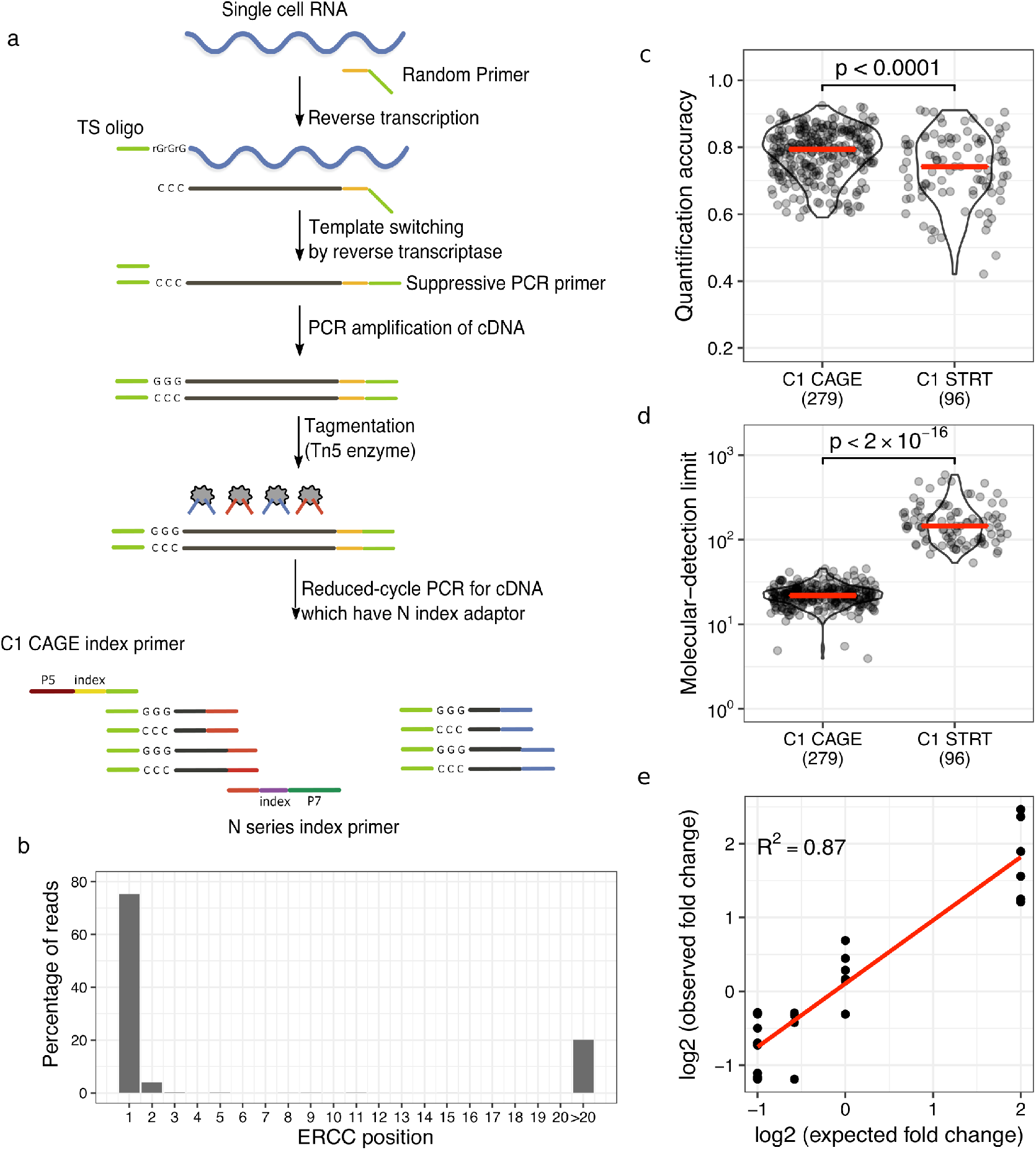
C1 CAGE method and performance. (a) Schematic of the C1 CAGE method. Tn5 enzymes are loaded with two different adaptors: N (red) and S (blue). P5, P7: Illumina sequencing adaptors. (b) Percentage of reads aligning to the 5’-end of ERCC spike-ins by nucleotide position. (c, d) Comparison between C1 CAGE and C1 STRT Seq (data from doi:10.1038/nmeth.4220). Red bars show median values. p-values from Welch two-sided Two Sample t-test shown. (c) Pearson correlation between expected and observed ERCC spike-in molecules. (d) The number of ERCC spike-in molecules required for a 50 % chance of detection. (e) Observed and expected fold-change ratios between ERCC mix1 and mix2. Linear regression line (red) and R-squared value shown.

To assess the specificity of 5’-end capture, we prepared libraries of A549 cells in the presence of synthetic “spike-in” RNAs, a set of 92 exogenous control transcripts with defined abundances developed by the External RNA Controls Consortium (ERCC)(Munro *et al*., 2014). We analyzed the positions of forward reads on these spike-ins and found that ∼80% of their 5’-ends align to the first base (Figure 1b), supporting the specificity of 5’-end capture in C1 CAGE. Of the remaining reads, about half of them can be explained by “strand-invasion” events, which are artefacts arising from interruption of first strand synthesis due to complementarity with the template switching oligonucleotide and can be identified based on the upstream sequence of the read(Tang *et al*., 2013). Next, we assessed the quantification accuracy and molecular detection limit(Svensson *et al*., 2017). For quantification accuracy, measured as the Pearson correlation between the input spike-in amounts and the observed read counts, C1 CAGE displayed a median of 0.79, slightly higher (Welch Two Sample t-test, two-sided: t=4, df=127.6, p < 0.0001) than C1 STRT Seq (median of 0.74, Figure 1c). For detection limit, measured as the median number of spike-in molecules required to give a 50% chance of detection, C1 CAGE displayed a median of 22, which is significantly more sensitive (Welch Two Sample t-test, two-sided: t=−14, df=94.2, p < 2.2e-16) compared with C1 STRT Seq (median of 146, Figure 1d). Finally, we assessed the ability of C1 CAGE to detect differential expression by comparing libraries prepared using two reference mixtures of spike-ins with fixed ratios of input amounts at 4, 1, 2/3 and 1/2 fold difference. Fitting a linear model we find an R-squared value of 87%(Figure 1e). These results demonstrate that C1 CAGE specifically captures the 5’-end of transcripts, has quantification accuracy and detection sensitivity comparable to other C1-system methods, and reliably detects differential expression with high accuracy.

### Color multiplexing

Taking advantage of the imaging capacities of the C1 system, we devised a strategy to multiplex samples within the same C1 CAGE replicate, by labelling cells with different Calcein AM dyes to encode sample information and monitor cell viability at the same time. Based on this approach, we multiplexed samples of A549 cells stimulated with TGF-β in a time-course at three time-points (0, 6, and 24 h, in triplicates) by permuting the Calcein AM dyes for each time point in each replicate (Figure 2a). The three C1 CAGE replicates were sequenced to a median depth of 2.4 million raw read pairs per cell. Analyzing the genomic distribution of forward read 5’-ends per replicate, a mean of 34% and 0.7% of reads were aligned to promoter and enhancer CAGE clusters, respectively (Figure 2b). Subsampling analysis demonstrates the number of CAGE clusters detected in most single-cells are saturated at the current sequencing depths, with a median of 2,788 CAGE clusters detected per cell (Figure 2c). To demultiplex time points, we localized the cells in their capture chambers on the IFCs and quantified their fluorescence in the red and green channels, identifying 40, 41 and 70 cells for time points 0, 6 and 24 h, respectively. Following the scran pipeline(Lun, McCarthy and Marioni, 2016) we removed 15 unreliable cells, arriving at the final set of 136 high quality cells. Initially, we observed a strong batch effect with principal components analysis (PCA), where cells cluster by replicate (Figure S1a). However, our experimental design ensured that each replicate contained cells for each time point, allowing us to correct for this batch effect using linear modelling. After batch correction cells were clustered by time points rather than by replicate (Figure S1b). After removing low abundance CAGE clusters, our final dataset detected 18,687 CAGE clusters, covering 9,809 GENCODE genes (Figure S2; annotation breakdown) and 826 FANTOM5 enhancers. For comparison, we generated corresponding bulk CAGE data using the nAnT-iCAGE method(Murata *et al*., 2014) for each sample (0, 6, and 24 h, in triplicates) sequenced to median a depth of 10.7M reads.

**Figure 2:**
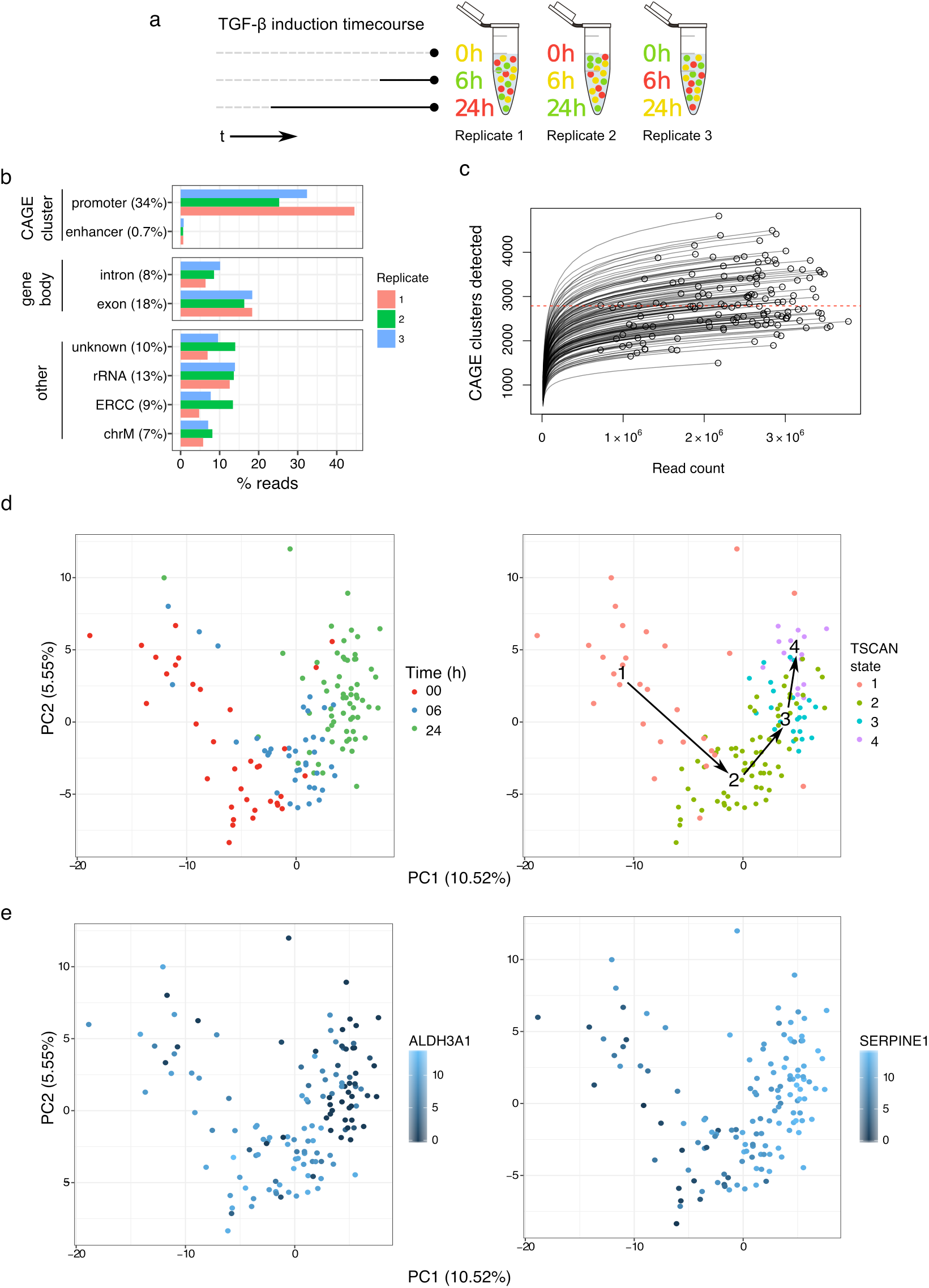
Multiplexing time course strategy. (a) Different color combinations of cells from each time point are added to each replicate. (b) Forward read 5’-end counts by annotation category. Mean read percentage per category shown in brackets. (c) Count of CAGE clusters within each cell after subsampling. Dashed red line at median (2,788). (d) PCA of cells performed on variable subset of CAGE clusters, percentage of variance explained by components shown, cells colored by time point and TSCAN state. (e) PCA of cells performed on variable subset of CAGE clusters, percentage of variance explained by components shown, cells colored by expression values for the marker genes *ALDH3A1* and *SERPINE1* demonstrating that the dynamics of TGF-β response are captured by the TSCAN states.

### Dynamic TSS regulation upon TGF-β treatment

To identify TSS that are dynamically regulated during TGF-β treatment, we performed pseudotime analysis on a variable subset of CAGE clusters with TSCAN(Ji and Ji, 2016). TSCAN divided the pseudotime ordering into four distinct states, which showed considerable consistency with the time points, as seen by PCA (Figure 2d). We also confirmed the consistency of the TSCAN states by visualizing the expression levels of two highly variable CAGE clusters for known EMT marker genes, *ALDH3A1* and *SERPINE1*, which showed a clear shift in expression levels from 0 h to 24 h (Figure 2e). To understand the influence of the cell cycle on how TSCAN defined the states, we calculated G2M scores with the cyclone package using the pre-calculated data trained on human embryonic stem cells (hESCs)(Scialdone *et al*., 2015; Leng *et al*., 2015). The clear separation of scores between states 1 and 2 points to the possibility that half (16/35) of 0 h cells were in proliferative states prior to TGF-β stimulation (Figure 2d and Figure S3).

To identify genes that are co-regulated across the TSCAN states, we performed Weighted Gene Co-Expression Network Analysis (WGCNA)(Langfelder and Horvath, 2008), correlating CAGE cluster expression levels across cells. We identified five co-expressed modules: Suppressed (n=1,041), Weak Responding I (n=825) & II (n=164), Early Responders (n=1,775), and Late Responders (n=2,223). We visualized their trajectories across the pseudotime using eigengene profiles to represent the average behavior and show two CAGE clusters from each module with eigengene correlation coefficient of at least 0.3 with p-value less than 0.1 (Figure 3a, b). The module labels were assigned based on these trajectory visualizations: Suppressed, Early and Late Responders represent those genes that undergo strong expression changes with TGF-β activation, whereas Weak Responding I and II represent those with little or no changes in their transcription.

**Figure 3:**
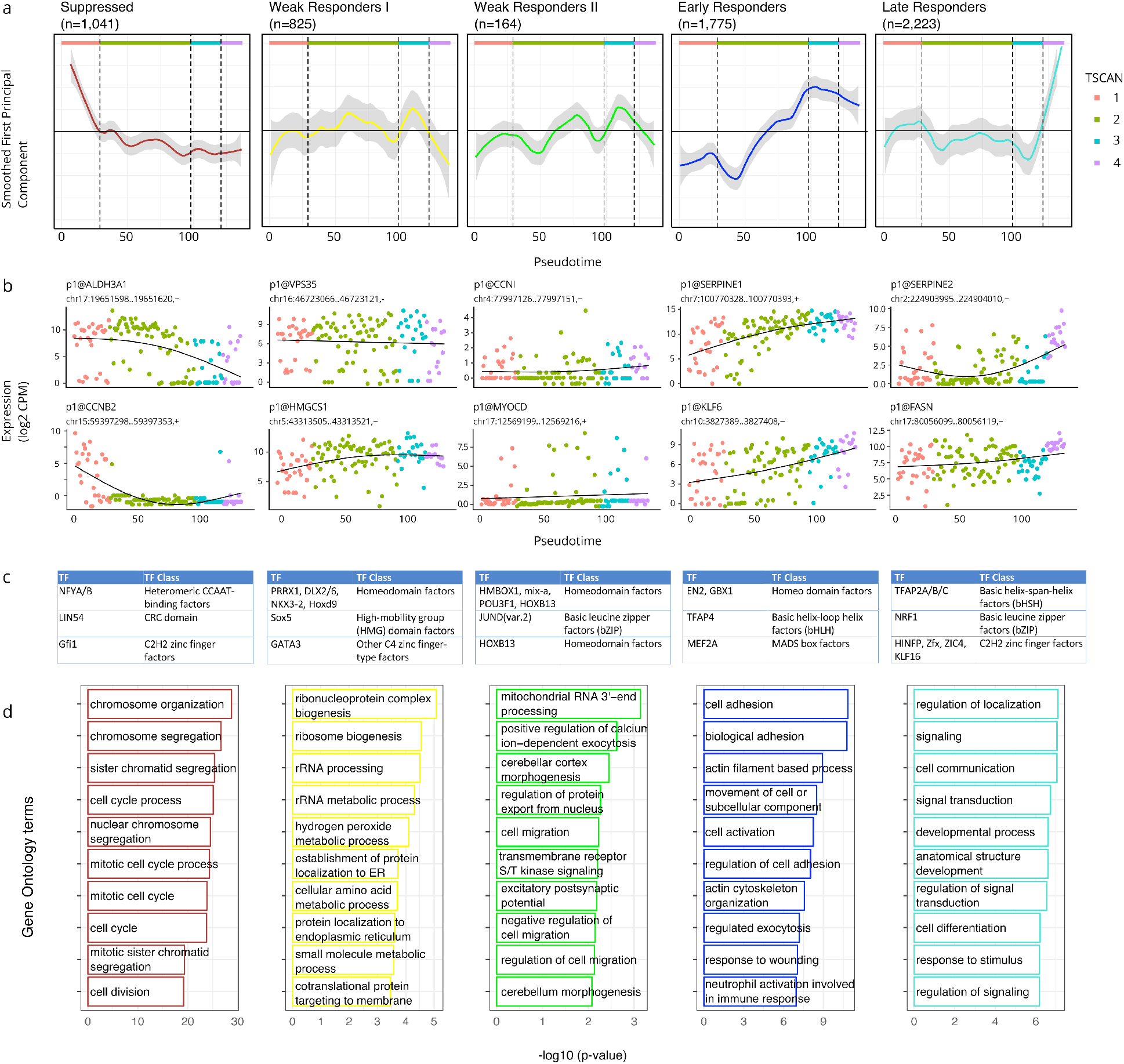
WGCNA clusters of response to TGFβ. (a) WGCNA results in 5 different modules, 3 of which show clear response behavior to TGF-β (Suppressed, Early Responders, Late Responders). (b) Example CAGE peaks from each module. (c) Top three enriched TF binding profiles in each module. (d) Functional analysis using edgeR’s implementation of GOseq. Top over-represented GO terms for biological processes are shown.

To understand the biological contexts of these modules, we investigated the enrichment of transcription factor binding motifs (Mathelier *et al*., 2016, Arenillas *et al*., 2016) and Gene Ontology (GO) terms in each module. Examining motifs enriched in all modules against a randomly generated GC-matched background, we find that the ETS-related factors are most prominent, such as ETVn, ETSn, ELKn, FLI and NFYx factors (Figure S4). The ETS family of transcription factors is well defined to promote metastasis progression in EMT process(Ell and Kang, 2013).

Examining each module individually against the combined background of all the other modules (Figure 3c, d) we observe the Suppressed Module enriched in GO terms related to DNA replication and the cell cycle. It has been reported that early after TGF-β treatment, the expression of multiple genes that play key roles in regulating cell cycle progression are suppressed(Schneider, Tarantola and Janshoff, 2011). We observe suppressed expression of *CCNB2* known to interact with the TGF-β pathway in promoting cell cycle arrest(Liu *et al*., 1999) and of *ALDH3A1* known to affect cell growth in A549 cells(Moreb *et al*., 2008). We also observe enriched motifs for the cell cycle regulators *LIN54* and GFI1(Basu *et al*., 2009; Sadasivam and DeCaprio, 2013). CAGE clusters in the Suppressed module are more highly expressed in TSCAN state 1, which may represent cells which have not yet fully undergone TGF-β induced G1 arrest as explained above.

Within the Early Responders and Late Responders modules we observe canonical TGF-β response genes, including *KLF6* known to suppress growth through TGF-β transactivation(Botella *et al*., 2009) and marker genes for EMT such as *SERPINE1* and *FASN*. TGF-β is one of the key signal transduction pathways leading to EMT and several lines of evidence implicate increased TGF-β signaling as a key effector of EMT in cancer progression and metastasis(Massagué, 2008; Ikushima and Miyazono, 2010; Heldin, Vanlandewijck and Moustakas, 2012). We observed upregulation of mesenchymal marker genes, with a clear increase in *Vimentin* (*VIM*) expression starting during TSCAN state 2, and expression of *N-cadherin* (*CDH2*) not detected until TSCAN state 2, and then expressed within a subset of cells(Figure S5).

Within the Late Responders module we observe enrichment for TFAP2 family transcription factors (TFs) (Figure 3c), suggesting that they might play a role in the late response to TGF-β signaling. We examined their expression profiles in both the single-cell and bulk data, and found *TFAP2C* to have a strong time-dependent expression profile in bulk data, and sporadic expression in TSCAN states 1 and 2 but not in the later states(Figure S6). *TFAP2C* is a known marker gene in breast cancer biology, its loss resulting in increased expression of mesenchymal markers associated with the transition from luminal to basal subtypes(Cyr *et al*., 2015) and the direct repression of cell cycle regulator CDKN1A(Williams *et al*., 2009; Wong *et al*., 2012).

Examining differences between the Early Responders and Late Responders modules, we find GO terms relating to cell adhesion enriched in Early Responders genes, and GO terms related to cell communication and signaling enriched in the Late Responders genes (Figure 3d).

To further dissect the functional heterogeneity in response to TGF-β, we revisited TSCAN states analysis and explored states 3 and 4 which we observe 24 h post stimulation (Figure 2d). To examine differences between the two states, we performed gene set enrichment analysis amongst CAGE clusters from the Early Responders and Late Responders modules with Camera(Wu and Smyth, 2012) and find a number of gene sets significantly upregulated in TSCAN state 4 including Epithelial to Mesenchymal transition (38 genes, FDR=0.003; full results in Supplementary Table 2). This suggests bi-phasic state in response to TGF-β 24 h post stimulation. Interestingly, a previous study implicated bi-phasic state with more severe morphological changes such as cell-to-cell contacts occurring from 10 to 30 h (Schneider, Tarantola and Janshoff, 2011). Thus, the additional states inferred from the pseudotime analysis reveal the asynchronous progression cells upon TGF-β treatment, which would not have been possible with bulk analyses of the three time points.

### eRNA in C1 CAGE

Next we asked whether C1 CAGE can detect the dynamic expression of eRNAs. We and others have reported that bidirectional transcription is associated with enhancer activity(Andersson *et al*., 2014). We observe a similar signature of bidirectional transcription at enhancers detected in pooled C1 CAGE and bulk CAGE data sets (Figure 4a), as well as a similar enrichment of DNase hypersensitivity and H3K27 acetylation, indicating that C1 CAGE unambiguously detected the transcription of eRNAs at these active enhancer regions (Figure 4b). To further examine the bidirectionality of eRNAs at a single-cell level, we selected enhancers with at least 10 reads in at least 5 cells to filter for the most widely and strongly detected enhancers and avoid bias due to dropout. For each enhancer, we calculated a bidirectionality score in pooled single-cells ranging from 0 to 1, with 0 being perfectly balanced bidirectional and 1 being perfectly unidirectional. Examining a set of enhancers (n=32) with balanced transcription, we calculated their bidirectionality score within single-cells, where these enhancers were unidirectionally transcribed (single-cell bidirectionality scores >0.9) (Figure 4c, shown in detail for one enhancer in Figure 4d), indicating that simultaneous transcription of eRNAs from both strands is generally not observed within single-cells.

**Figure 4:**
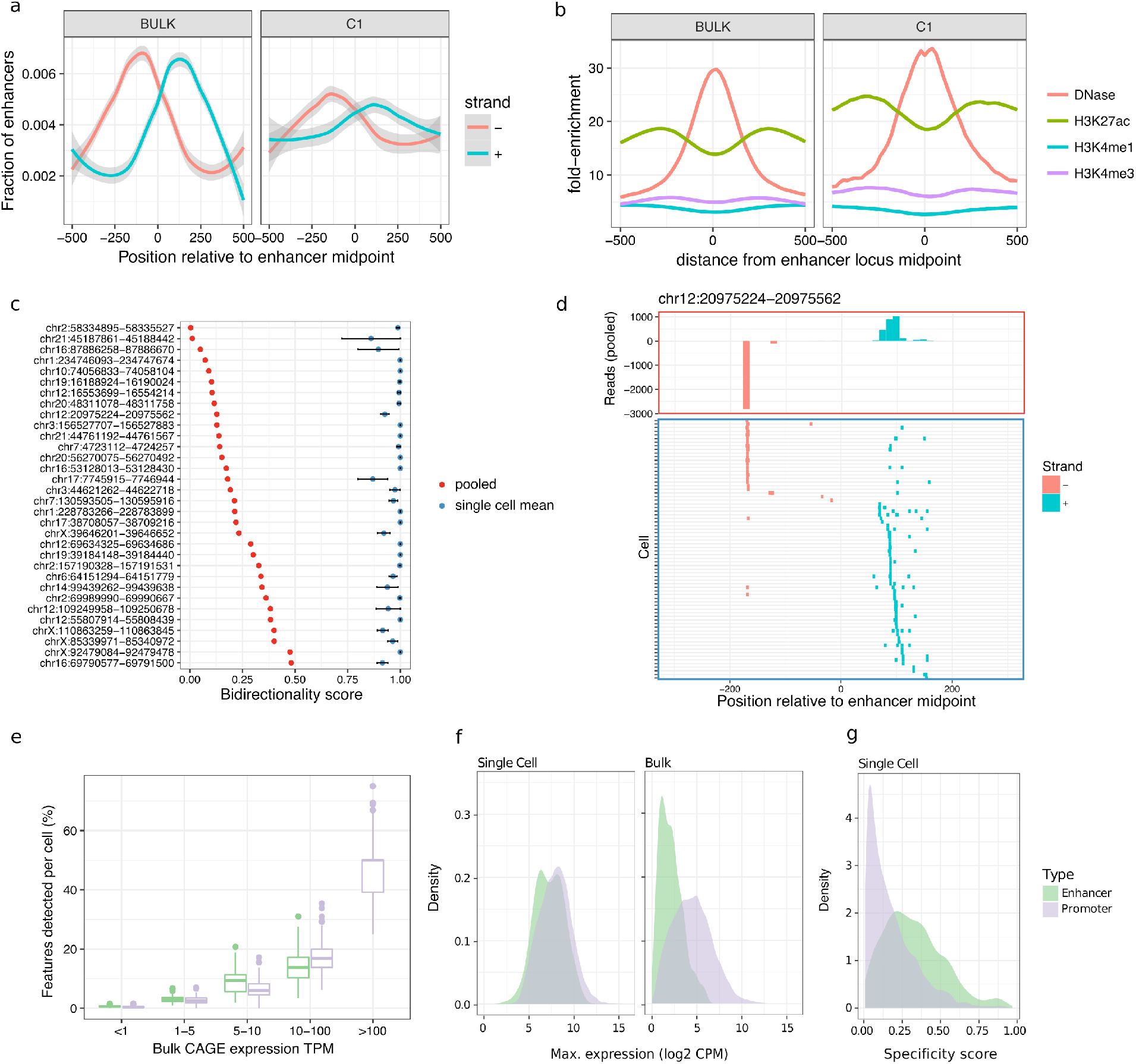
Enhancer analysis at single-cell resolution. Comparison of enhancers detected by bulk CAGE and pooled C1 CAGE data (a) showing bidirectional read profiles smoothed by generalized additive model and (b) epigenetic profiles. (c) Bidirectionality analysis scores (0: equally bidirectional; 1: fully unidirectional) at selected enhancers for pooled cells (red dots) and single-cells (blue dots: mean; black bars: standard error). (d) Example locus on chromosome 12: read profile histogram (upper box), and read presence or absence in single-cells (lower box). (e, f, g) Comparison of enhancers and gene promoters in C1 CAGE and bulk CAGE: (e) Fraction of bulk features detected within each cell, stratified by bulk expression level, (f) Density plots of the maximum expression levels, (g) Specificity score distribution in single-cell data. Lower scores: broad expression (expressed in more cells); higher scores: more specific/enriched expression (fewer cells).

Although most enhancers were sporadically detected among single-cells, they were detected at a similar level to promoters in single-cells when controlling for expression level (Figure 4e). To assess if enhancers are generally lowly expressed among cells or if they are highly expressed in a subset of cells, we compared the distributions of the maximum expression levels of enhancers and promoters within single-cells and in the bulk data sets (Figure 4f). While the expression of enhancers is generally lower than that of promoters in the bulk data sets, they have similar distributions of expression levels within single-cells. To further evaluate the specificity of enhancer expression in single-cells, we devised a specificity score ranging from 0 to 1, with 0 being ubiquitously expressed (i.e. broad expression in many cells), and 1 being specifically expressed (i.e. expression restricted to few cells). We found that enhancers show significantly higher specificity scores than promoters (Figure 4g; Kolmogorov-Smirnov test, D=0.36562, p-value<2.2e-16). This suggests that enhancers behave similarly to promoters which are expressed in transcriptional bursts(Suter *et al*., 2011; Bahar Halpern *et al*., 2015) but have fewer numbers of cells where bursts of expression take place, which in turn are averaged out by the total population of cells used to obtain the bulk RNA profile.

### FISH validation

To validate the ability of C1 CAGE to detect eRNAs in single-cells, we used smFISH(Femino *et al*., 1998; Raj *et al*., 2008) to visualize the expression of these transcripts through the TGF-β time course in A549 cells. We first selected intergenic enhancers, filtering out those that overlapped any known transcript models in GENCODEv25, and ranked them by their expression levels. We then searched for their proximal promoters within the same topologically associated domain (TAD) as the potential targets of these enhancers. We selected three enhancers, two of which displayed expression changes across the time-course (Figure S6, S7) and were adjacent to genes known to be involved in TGF-β response, *KLF6* and *PMEPA1* (*KLF6*-eRNA1 at chr10:3929991-3930887 and *PMEPA1*-eRNA1 at chr20:56293544-56293843, respectively), and a third enhancer (*PDK2*-eRNA1 at chr17:48105016-48105270) adjacent to *PDK2*.

In line with previous reports(Rahman *et al*., 2016; Shibayama, Fanucchi and Mhlanga, 2017), smFISH for eRNAs gave rise to punctate spots mainly restricted to the nuclei and always no greater than the copy number of the chromosome harboring the enhancer, suggesting that these eRNAs are expressed in low-copy-number and remain at or near their site of transcription. Targeting eRNAs on both strands with the same color, smFISH displayed expression profiles similar to C1 CAGE for the *KLF6*-eRNA1 and *PMEPA1*-eRNA1 enhancers that were upregulated in the C1 CAGE time-course data (Figure 5a, b). In contrast, PDK2-eRNA1, whose expression remained steady in smFISH, decreased in the number of cells with signal across the time course in C1 CAGE (Figure S8a).

**Figure 5:**
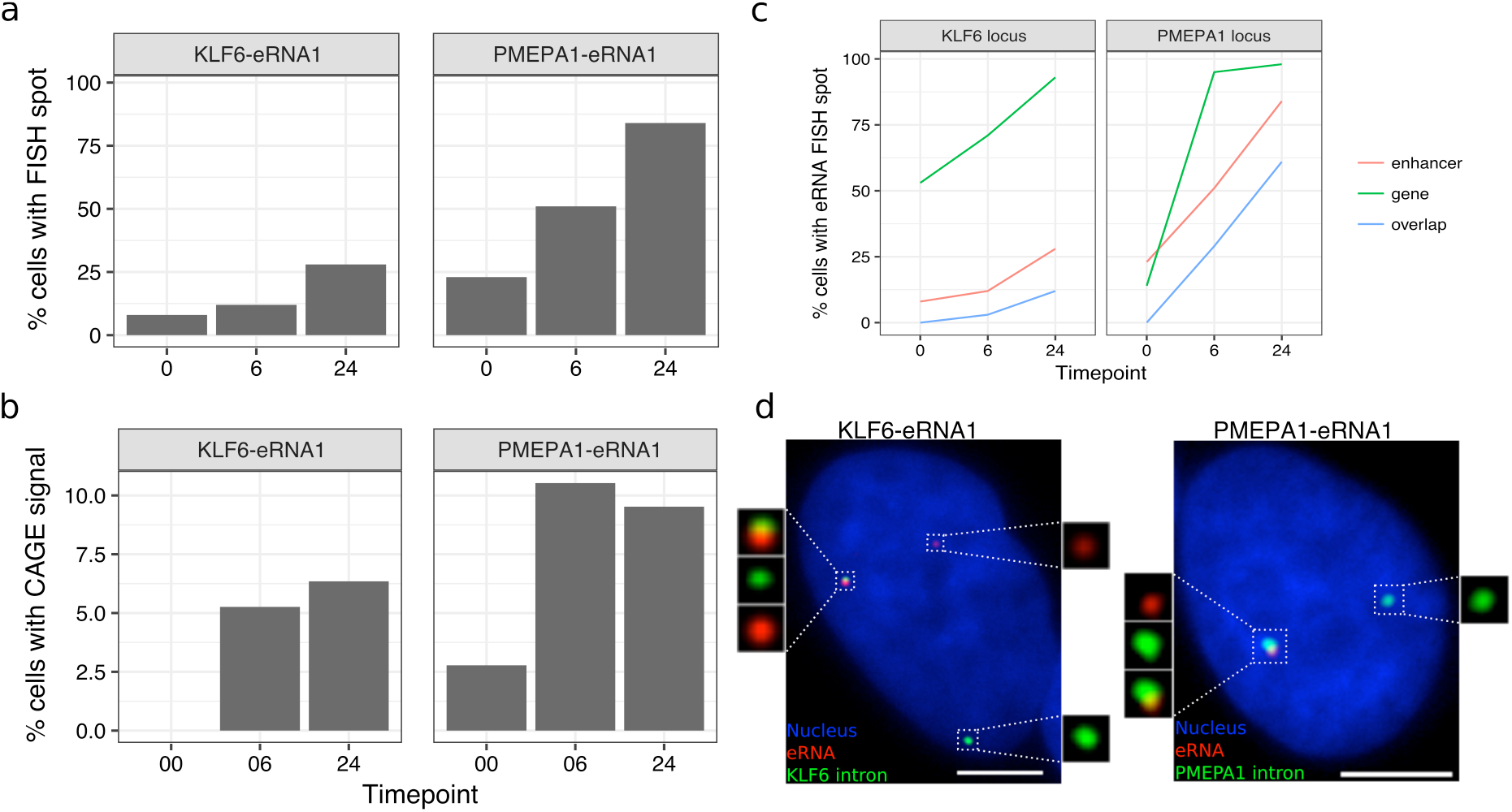
Enhancer and promoter profiles in smFISH. (a, b) Proportion of cells with KLF6-eRNA1 and PMEPA1-eRNA1 detected by (a) FISH, (b) C1 CAGE. (c) Proportion of cells with detected gene intron, enhancer locus and cells with spot overlap at the KLF6 and PMEPA1 loci. (d) Representative images showing gene intron and enhancer locus detection by FISH. Bar = 5 μm. n=100 per time point.

For validation of our findings that eRNA were expressed unidirectionally within single-cells, we also targeted the + and − strands of the *KLF6*-eRNA1 and *PMEPA1*-eRNA1 eRNAs in separate colors. In agreement with the C1 CAGE data for these particular enhancers, the majority of the detected spots belonged to eRNAs from only one strand (Figure S8b). In nuclei where eRNAs from both strands were detected, spot co-localization was rare, confirming our suggestion that simultaneous bidirectional transcription of enhancers from single alleles is a rare event.

Next, we checked for the association of eRNAs with the transcription of nearby genes using smFISH. Visualization of nearby gene transcription was achieved by targeting only the intronic portion (i.e. nascent RNA). Colocalization of an enhancer RNA spot with a nascent RNA spot would suggest the presence of the enhancer RNA at the site of gene transcription, potentially implicating the enhancer’s role in promoter activity. Interestingly, nascent transcription of nearby protein coding genes showed similar expression kinetics to the enhancers themselves indicated by increased co-expression of both the protein coding gene and the nearby eRNA in TGF-β stimulated cells (Figure 5c, d, S8c). For KLF6-eRNA1 and PMEPA1-eRNA1, we observed time-dependent increase in colocalization and in the number of nuclei with colocalized spots (Figure 5c, d, S9c). In unstimulated cells displaying a basal level of expression of both enhancer and promoter, colocalization of spots could not be observed. This suggests a stimulus-dependent co-activation of enhancer and its association with the nearby promoter. However, a significant portion of transcription sites expressed no enhancer RNA. Possible reasons include a potential delayed interval between transcription events from an enhancer and promoter, during which most enhancer RNA is rapidly degraded. It is also possible that other nearby enhancers may exert their effect on a target promoter. In summary, smFISH could validate enhancer expression, including strand specificity, in single-cells as detected by C1 CAGE.

## Discussion

We examined the response to TGF-β in A549 cells to uncover dynamically regulated promoters and enhancers at single-cell resolution. We highlight enhancer dynamics at single-cell resolution and suggest transcriptional bursting of enhancers, and that while enhancers show bidirectional eRNA transcription in pooled cells, transcripts are generally mutually exclusive.

Among the eight publicly available transcriptome methods for the C1 platform (Supplemental table 1), only C1 CAGE provides strand-specific whole-transcriptome coverage: its detection of 5’-ends is independent from transcript length and polyadenylation owing to the use of random primers. To make the method more accessible, we used a commercially available tagmentation kit in which the transposase is loaded with two different adapters. This adaptation leads to half of the tagmentation products being lost in the process of library preparation. The use of custom loaded transposase, such as in C1 STRT Seq(Islam *et al*., 2014), would allow reduction of the final PCR amplification by one cycle and enrich extracted reads in the sequencing library, however at the expense of not using standard reagents.

C1 CAGE has single-nucleotide resolution of transcript 5’-ends, as demonstrated by the data on ERCC spike-ins, where 80% of read one 5’-ends align to the first base. In this study, we did not use ERCC spike-ins for normalization of endogenous genes, preferring to use size factors computed from pools of cells(Lun, Bach and Marioni, 2016), as experimental noise due to spike-in preparation may be introduced(Svensson *et al*., 2017). Notably, we could detect the ERCC spike-ins even if they are not capped. Nevertheless, C1 CAGE shows a preference for capped ends, as suggested by the fact that the C1 CAGE library contained only 13% reads from ribosomal RNAs. While this range of ribosomal RNA is acceptable, further reduction might be achieved through the use of pseudo-random primers(Arnaud *et al*., 2016).

The template-switching oligonucleotides (TSOs) included Unique Molecular Identifier (UMIs)(Islam *et al*., 2014), however we have not utilized them for molecular counting, because the TSOs carried over from the reverse-transcription could prime the subsequent PCR reaction while tolerating mismatches on the UMI sequence, thus causing a high level of mutation rate (as evidenced by the fact that most UMIs are seen only once). Nevertheless, PCR duplicates are partially removed from our data due to the use of paired-end sequencing, as our alignment workflow collapses the pairs that have exactly the same alignment coordinates. Further improvements of the C1 CAGE might address the mutation rate in UMIs. However, attempts to make the TSOs heat-labile by using full RNA composition have not been successful so far (CP and SK, personal communication).

Batch effect is a common problem in single-cell RNA-seq, and failing to account for this can lead to cofounding biological interpretations. We introduced, for the first time, an image based approach to decode multiplex samples by using two colors of Calcein AM and their combinations. Moreover, the platform further allows the usage of a larger number of colors or alternatives to Calceins, such as MTT, ATP or MitoBright, which are generally used for live cell monitoring. For instance, we previously used FUCCI fluorescent reporters to detect cell cycle phases(Böttcher *et al*., 2016). Other potential applications could include the detection of cytoplasmic or nuclear localizations of fluorescent-labelled transcription factors, or cell division counting with fluorescent probes.

Our cell cycle classification was performed using a model trained on data from H1 hESCs expressing the cell-cycle indicator FUCCI in the C1 system(Leng *et al*., 2015). While training data from phased A549 single-cells would have been preferable, models trained on mouse ESC have also been applied to other cell types with accuracy(Scialdone *et al*., 2015). However, because the hESC training data was obtained from a 3’-end capture protocol, it may contain different experimental biases that are distinct from our C1 CAGE method. Therefore, these results should be interpreted with caution, and we did not exclude cells based on this classification.

The chemistry implemented in C1 CAGE–template switching, random priming, and interrogation of 5’-ends–revealed promoter and enhancer activities in lung adenocarcinoma cell line. Enhancers have previously been defined by a signature of balanced bidirectional transcription in bulk data(Andersson *et al*., 2014). Here we suggest that this signature arises due to generally mutually exclusive transcription from each strand within single-cells. We also suggest for the first time that while eRNAs appear lowly expressed in bulk data, they can be expressed at similar levels to gene promoters within single-cells, although they are expressed in a more restricted subset of cells—i.e. displaying transcriptional bursting.

Notably, C1 CAGE is not restricted to the use in the C1 platform. Indeed, some of the changes introduced in C1 CAGE are also available for bulk nanoCAGE libraries in our latest update(Poulain *et al*., 2017). Moreover, the C1 CAGE chemistry might be applicable to profile large numbers of single-cells with droplet based single-cell capture methods. Droplet technologies are more robust to variations of the cell size, and have higher throughput, although they do not allow for the association of imaging. Five-prime-focused atlases will yield greater insights towards promoter and enhancer activities in various biological systems.

## Online Methods

### Cell culture and TGF-β stimulation

A549 cells (ATCC CCL 185) were grown at 37 °C with 5 % CO_2_ in DMEM (Wako, Lot: AWG7009) with 10 % fetal bovine serum (Nichirei Bioscience, Lot 1495557) and penicillin/streptomycin (Wako, Lot 168-23191). At 0 h, 10^6^ cells were seeded in 10 cm dishes (TRP, Cat. num. 93100). At 24 h, the medium was replaced with DMEM without serum after 3 times washing with PBS (Wako, Lot 045-29795). At 48 h, one third of the dishes were stimulated by treating with 5 ng/ml TGF-β (R&D systems, USA, Accession #P01137). At 66 h, the second third was stimulated with the same treatment. At 72 h, cells for each treatment duration (0 h, 6 h 24 h) were collected and stained with combinations of Calcein AM and Calcein red-orange, (Thermo Fisher Scientific, L3224 and C34851). Transcriptome alignment of the C1 positive controls against 79 reference genomes of Mycoplasma or Acholeplasma, including Mycoplasma hominis, confirmed the absence of contamination.

### Cell capture

Calcein stained cells were captured in C1 Single-cell Auto Prep Integrated Fluidic Circuits (IFC) for mRNA Seq, designed for medium-sized (10 to 17 μm) cells (Cat. Num. 100-5760), following manufacturer’s instructions (PN 100-7168). In brief, 60 μl of 2.5 × 10^5^ cell/ml and 40 μl C1 suspension buffer were mixed (all C1 reagents were from Fluidigm), and 20 μl of this mix was loaded into a primed IFC, and processed the script “mRNA Seq: Cell load (1772x/1773x)”

### Imaging

After loading, IFCs were imaged on INCell Analyzer 6000 (GE Healthcare). Calcein AM was excited at 488 nm and imaged with a FITC fluorescence filter (Semrock). For Calcein red-orange, excitation was at 561 nm (TexasRed; Semrock). Eleven focal planes per chamber and channel were acquired and manually curated to detect empty, dead, singlet, doublet or multiplet cells in the capture site. In case of single-plane imaging, we used the Cellomics platform like in Böttcher et al., 2016^42^ (with a green filter (excitation bandwidth: 480-495 nm, emission bandwidth: 510-545 nm), and with a red filter (excitation bandwidth: 565-580 nm, emission bandwidth: 610-670 nm (Thermo Scientific)). Processed and raw single-cell images are available for download from http://single-cell.clst.riken.jp/riken_data/A549_TGF_summary_view.php

### Lysis, reverse transcription and PCR for C1-CAGE

Single-cell RNA extraction and cDNA amplification were performed on the C1 IFCs following the C1 CAGE procedure that we deposited in Fluidigm’s Script Hub. (https://www.fluidigm.com/c1openapp/scripthub/script/2015-07/c1-cage-1436761405138-3). In brief, cells were loaded in lysis buffer (C1 loading reagent, 0.2 % Triton X, 15.2 U Recombinant Ribonuclease Inhibitor, 37.5 pmol reverse-transcription primer, DNA suspension buffer, ERCC RNA Spike-In Mix I or II (Thermo Fisher, 4456653) diluted either 20,000 times (protocol revision B) or 200 times (revision A)), and lysed by heat (72 °C 3 min, 4 °C 10 min, 25 °C 1 min). First-strand cDNAs were reverse transcribed (22 °C 10 min, 42 °C 90 min, 75 °C 15 min) in C1 loading reagent, First Strand buffer, 0.24 pmol dithiothreitol, 15.4 nmol dNTP Mix, betaine, 24.8 U Recombinant Ribonuclease Inhibitor, 175 pmol template-switching oligonucleotide, and 490 U SuperScript III. The cDNAs were amplified by PCR (95 °C 1 min, 30 cycles of 95 °C 15 s, 65 °C 30 s and 68 °C 6 min, 72 °C 10 min) in a mixture containing C1 loading reagent, PCR water, Advantage2 PCR buffer (not SA), dNTP Mix (10 mM each), 24 pmol PCR primer, 50 × Advantage2 Polymerase Mix. The PCR products (13 μl) were then harvested in a 96-well plate and quantified with the PicoGreen (Thermo Fisher, P11496) method following the instructions from Fluidigm’s C1 mRNA-Seq protocol (PN 100-7168 I1). On-chip cDNA amplification with 30 PCR cycles yielded 1.0 ng/μl in average from single cell. A subset of the samples were further controlled by size profiling on the Agilent Bioanalyzer with High Sensitivity DNA Chip.

### Tagmentation reaction, index PCR and sequence

Amplified cDNAs were diluted to approximately 0.2 ng/μl following the C1 mRNA-Seq protocol, fragmented and barcoded by “tagmentation” using the Nextera XT kit (Illumina, cat. num. FC-131-1096-RN) following the instructions from Fluidigm’s C1 mRNA-Seq protocol (PN 100-7168 I1), except that we used custom forward PCR primers (dir#501-508/N701-N712, Supplementary Table 3). The final purified library was quality-controlled on a High-Sensitivity DNA Chip and quantified with the KAPA Quantification Kit (Nippon Genetics). Nine pmol were sequenced and demultiplexed on Illumina HiSeq 2500 High output mode (50 nt paired end).

### CAGE processing

In forward read (Read 1) sequences, linkers were removed and unique molecular identifiers were extracted using TagDust2(Lassmann, 2015). Reverse read (Read 2) sequences were then filtered with the program syncpairs (https://github.com/mmendez12/sync_paired_end_reads) to restore the pairing. The pairs were then filtered against the sequences of the human ribosomal RNA locus (GenBank ID U13369.1), and linker oligonucleotides using TagDust2 v2.13 in paired-end mode. They were then aligned to the human genome version hg19 with Burrows Wheeler Aligner (BWA)’s “sampe” method(Li and Durbin, 2010) with a maximum insert size of 2,000,000. To map the reads on the ERCC spikes at a single nucleotide resolution, we prepared reference sequences of the T7 transcription of the ERCC plasmids, which are now available from the NIST’s website (https://www-s.nist.gov/srmors/certificates/documents/SRM2374_putative_T7_products_NoPolyA_v1.fasta) (many RNA-seq studies previously published aligned their reads only to the sequence of the plasmid inserts, which lack transcribed linker sequences, which are essential for aligning CAGE reads precisely to the 5’ ends). The properly aligned pairs were then converted to BED12 format with the program pairedBamToBed12 (https://github.com/Population-Transcriptomics/pairedBamToBed12) with the option “-extraG”, and assembled in CAGEscan fragments with the program umicountFP (https://github.com/mmendez12/umicount/). This workflow was implemented in the Moirai system (PMID:24884663) and a prototype implemented in a Jupyter notebook is available on GitHub (https://github.com/Population-Transcriptomics/C1-CAGE-preview/blob/master/OP-WORKFLOW-CAGEscan-short-reads-v2.0.ipynb). The 5’ ends of the CAGEscan fragments represent TSS in the sense of Sequence Ontology’s term SO:0000315 (“The first base where RNA polymerase begins to synthesize the RNA transcript”).

### Bulk CAGE

Bulk CAGE data was generated by nAnT-iCAGE method(Murata *et al*., 2014). Briefly, 5 μg of total RNA prepared from remaining A549 cells after C1 loading. cDNA was reverse transcribed using SuperScript III reverse transcriptase, biotinylated and cap trapped to capture 5’ completed cDNAs. Each cDNAs were barcoded and purified. Libraries were sequenced on Illumina HiSeq 2500 High output mode (50 nt single read).

### Image curation and time point demultiplexing

We used the Bioconductor package CONFESS (LOW D and MOTAKIS E (2017). *CONFESS: Cell OrderiNg by FluorEScence Signal*. R package version 1.6.0) to detect the cells present in the capture chambers, and quantify the fluorescence in the Green and Red channels. In addition, two curators visually screened the images to confirm the presence of cells, and to detect doublets when focal stacks were available. The final annotation reflects the consensus of the three curations. The results were then cross-checked with other quality control parameters, in particular the amount of cDNAs yielded by the C1 runs, and the fraction of spikes and ribosomal RNA in the libraries. In case of conflicting results, chamber images were re-inspected and re-annotated, if necessary.

### ERCC spike-in analysis

Accuracy and molecular detection limits were calculated as in Svensson 2017(Svensson *et al*., 2017): The amount of input spike-in molecules for each spike, for each sample, in each experiment was calculated from the final concentration of ERCC spike-in mix in the sample. The calculation of the accuracy of an individual sample was determined with the Pearson correlation between input concentration of the spike-ins and the measured expression values. Molecular detection limit was calculated using the R function glm from the stats package.

### Read Annotation

The annotation used combined FANTOM5 robust cage clusters for promoters (http://fantom.gsc.riken.jp/5/datafiles/latest/extra/CAGE_peaks/) and enhancers (http://fantom.gsc.riken.jp/5/datafiles/latest/extra/Enhancers/). Promoter clusters were subtracted from enhancer clusters and annotated to their nearest GENCODEv25 within 500 bp where possible. A mask was added to remove rRNA, tRNA, small RNAs, unannotated promoters.

### Data Processing

After removing low quality cells and multiple single cells captured sites based on imaging data (SCPortalen)(Abugessaisa *et al*., 2018), the CAGE reads from the remaining 151 cells that overlapped the annotation CAGE clusters were summed together to create the raw counts matrix. This matrix was processed with the scran package(Lun, McCarthy and Marioni, 2016) version 1.6.6 in R 3.4.3 for quality control, filtering and normalization. Following the guideline suggested by the authors of scran, we first removed from our analysis 15 cells with 1) library sizes or feature sizes 3 median absolute deviations (MADs) below their median, or 2) mitochondrial proportion or spike proportion 3 MADs above their median, leaving us with 136 cells. All the cells that were dropped due to high spike proportion also had low library sizes and feature counts, whereas this was not necessarily true for those that were dropped due to high mitochondrial proportion. 14 out of the 15 removed cells were from the same C1 run (library 2), but there was no noticeable bias towards any particular time point (5, 3, 7 cells from 0h, 6 h, 24 h, respectively). We calculated the cell cycle phase scores using the cyclone method(Scialdone *et al*., 2015) for each cell. We filtered out low abundance features that were expressed in less than 2 cells or average counts of less than 0.3, leaving us with 18,687 features, of which 826 are FANTOM5 enhancers. These features were normalized with size factors calculated based on clusters of cells with minimum size of 30. We then performed mean-variance trend fitting using the whole endogenous feature set, building the sample replicate and Calcein staining variables into the model. We normalized the expression scores to correct for differences of sequencing depth, using a pooling-deconvolution approach(Lun, Bach and Marioni, 2016). We then detrended the data for possible C1 run and Calcein color effects. Lastly, we denoised the data by removing low-rank principal components. To produce the final normalized expression levels for downstream analyses, we reduced the technical noise using scran’s denoisePCA function based on the fitted data, then performed batch effect removal with the replicate and the Calcein stain as the covariates using limma package’s removeBatchEffect function. We selected high variance CAGE clusters (HVCs) as those with biological variation above the 75% quantile and false discovery rate less than 0.05 after decomposing the total variance for each gene into its biological and technical components using trendVar (scran). We also calculated the pairwise correlations among the HVCs and marked those with FDR greater than 0.05 as significantly correlating HVGs.

To create the pseudotime ordering with TSCAN (version 1.16.0), we selected the input feature set as the union of the significantly correlating HVCs, the top 100 HVCs and SC3(Kiselev *et al*., 2017) defined marker genes, totaling 290 CAGE clusters.

### WGCNA

WGCNA version 1.61 was used, with cut height detection threshold of 0.995, minimum module size of 100, signed network type, and merge cut height of 0.25. To reduce noise, we restricted ourselves to those features with mean expression greater than the median of the mean expression across all samples, and biological variation greater than the median. Also, to avoid having the same gene appearing in multiple clusters due to different promoters of the same gene being assigned as such, we only included the major promoter (highest sum of normalized expression across all samples) in the input set, which left us with 6,028 CAGE clusters as the input set.

### Motif analysis

Motif analysis was performed using CAGEd-oPOSSUM, which employs two separate scoring systems based on JASPAR 2016 transcription factor binding profiles, searching 500bp either side of CAGE clusters: 1) Z-scores, which counts the total number of a given motif found in the input set, and 2) Fisher score, which counts the number of input regions with the given motif. JASPAR motifs with information content greater than 8 bits were searched.

### Functional analysis

To see if we could identify any functional characteristics of the genes in each module, we performed a test of gene ontology term over-representation test using the edgeR’s goana function, which is an implementation of GOseq(Young *et al*., 2010). For input, we included those CAGE clusters that showed correlation coefficient of greater than 0.2 with p-value less than 0.1 with each module’s eigengene.

Camera gene set enrichment analysis(Wu and Smyth, 2012) was performed testing for differential expression between TSCAN states 3 and 4. For the input expression table, we selected the CAGE clusters that were included in the WGCNA analysis and were annotated with Entrezgene IDs. For the test set, we selected those CAGE clusters that showed correlation coefficient of greater than 0.2 with p-value less than 0.1 their module’s eigengene from the Early Responders and Late Responders modules. MSigDB Hallmark gene sets were used.(Liberzon *et al*., 2015)

### TADs

Out of 826 enhancers, 692 could be assigned to a topological association domain (TAD) identified in A549 cells from ENCODE Dataset GSE105600

### FISH

enhancer RNA lengths were estimated from the ENCODE A549 RNA-seq signal(Dunham *et al*., 2012). We designed oligonucleotide probes consisting of 20 nt targeting sequence using the Stellaris Probe Designer (Biosearch Tech). These sequences were flanked on both ends by 30 nt “readout sequence” serving as annealing sites for secondary probes that are labeled with a fluorescent dye(Chen *et al*., 2015). For each set of probes, all flanking sequences were identical, both on the 5’ and 3’ ends (Probes listed in Supplementary Table 4). Positive strand eRNA, negative strand eRNA and introns from each locus were assigned different flanking sequences to allow multiplexing. Secondary probes were labeled with either Atto 647 or Cy3 on the 3’ end. All probe sequences are listed in supplementary table 4. Briefly, cells were seeded onto coverslips overnight and were fixed in 4% formaldehyde in PBS for 10 min at room temperature.

After fixation, the coverslips were treated twice with ice-cold 0.1% sodium borohydride for 5 min at 4°C. Following three washes in PBS, the coverslips were treated with 0.5% Triton X-100 in PBS for 10 min at room temperature to permeabilize the cells. The coverslips were washed three times in PBS and treated with 70% formamide in 2x SSC for 10 min at room temperature, followed by two washes in ice-cold PBS and another wash in ice-cold 2x SSC. The coverslips were stored at 4°C for no longer than a few hours prior to hybridization. For hybridization, coverslips were incubated in hybridization buffer containing 252 nM primary probes overnight at 37°C inside a humid chamber. Hybridization buffer consisted of 10% formamide, 10% dextran sulfate, 2X SSC, 1μg/μl yeast tRNA, 2mM vanadyl ribonucleoside complex, 0.02% BSA. To remove excess probe, coverslips were washed twice in wash buffer made of 30% formamide, 2x SSC, 0.1% Triton X-100 for 30 min at room temperature and rinsed once in 2x SSC. For hybridization with secondary probes labeled with fluorescent dyes, coverslips were incubated in minimal hybridization buffer (10% formamide, 10% dextran sulfate, 2x SSC) containing 30 nM secondary probes for 3 h at 37°C inside a humid chamber. Coverslips were again washed twice in wash buffer for 30 min at room temperature and rinsed once in 2x SSC. Coverslips were mounted on glass slides using ProLong Gold Antifade Mountant with DAPI (Invitrogen). Imaging was done on a DeltaVision Elite microscope (GE) equipped with a sCMOS camera. Image processing and analysis were done using FIJI.

### Enhancer Analysis

For bidirectionality and epigenetic marks analysis a set of enhancers was selected overlapping ReMap(Chèneby *et al*., 2018) EP300 A549 binding sites. DNase, H3K27ac, H3K4me1 and H3K4me3 bigwig files were downloaded from the NIH roadmap epigenomics project(Roadmap Epigenomics Consortium *et al*., 2015) and processed with computeMatrix scale-regions from the deeptools package(Ramírez *et al*., 2016) for enhancer regions. Bidirectional enhancers were selected with at least 10 reads in at least 5 cells and a bidirectionality statistic was calculated as: *abs(plus strand reads - minus strand reads) / sum(reads)* ranging from 0 to 1 with 0 being equally bidirectional and 1 being fully unidirectional. 32 enhancers were selected with absolute score ≤ 0.5. This score was then calculated within each individual cell for these enhancers. The specificity score to indicate how broadly/specifically TSS were expressed we calculated Enrichment = *Max. Expression* / ∑(*Expression across all samples*).

### Data Availability

C1 CAGE sequence data from this study have been submitted to DDBJ (Project ID: PRJDB5282, Sample ID: SAMD00066188 - SAMD00066475). Alignments were uploaded to the ZENBU genome browser (Severin et al, 2014, PMID 24727769) and a default view is available at http://fantom.gsc.riken.jp/zenbu/gLyphs/#config=NMT9yTLnH59gIVssI9WRfD. In these two submissions the libraries numbered 1, 2 and 3 in this manuscript are numbered 4, 5 and 6, respectively, for historical reasons.

### Code Availability

Code used in this study is available at https://github.com/Population-Transcriptomics/C1-CAGE-manuscript

## Acknowledgements

This work was supported by a Research Grant from the Japanese Ministry of Education, Culture, Sports, Science and Technology (MEXT) to the RIKEN Center for Life Science Technologies. The authors wish to acknowledge RIKEN GeNAS for the sequencing of the libraries, and Fumi Hori for data deposition to DDBJ.

## Author Contributions

Conceptualization: TL, EA, CP, JWS

Ideas; formulation or evolution of overarching research goals and aims.

Data curation: TKo, AK, YH, MM, JSe, IA, CP

Management activities to annotate (produce metadata), scrub data and maintain research data (including software code, where it is necessary for interpreting the data itself) for initial use and later re-use.

Formal analysis: JM, AK, YH, EM

Application of statistical, mathematical, computational, or other formal techniques to analyze or synthesize study data.

Funding acquisition: PC, JWS

Acquisition of the financial support for the project leading to this publication.

Investigation: TKo, YS, SK, MB

Conducting a research and investigation process, specifically performing the experiments, or data/evidence collection.

Methodology: TKo, YS, SK, JL, CP, JWS

Development or design of methodology; creation of models.

Project administration: PC, CP, JWS

Management and coordination responsibility for the research activity planning and execution. Resources: ST, TA, MF, NR, JW, HS

Provision of study materials, reagents, materials, patients, laboratory samples, animals, instrumentation, computing resources, or other analysis tools.

Software: JM, AK, MB, MM, JSe, IA, AH, TL, CP

Programming, software development, designing computer programs implementation of the computer code and supporting algorithms, testing of existing code components.

Supervision: HS, TKa, TL, CCH, EA, CP, JWS

Oversight and leadership responsibility for the research activity planning and execution, including mentorship external to the core team.

Validation: TKo, YS

Verification, whether as a part of the activity or separate, of the overall replication/reproducibility of results/experiments and other research outputs.

Visualization: JM, AK, IA, CP

Preparation, creation and/or presentation of the published work, specifically visualization/data presentation.

Writing – original draft: TKo, JM, AK, YS, EA, CP, JWS

Preparation, creation and/or presentation of the published work, specifically writing the initial draft (including substantive translation).

Writing – review & editing: JM, CP, JWS

Preparation, creation and/or presentation of the published work by those from the original research group, specifically critical review, commentary or revision– including pre- or post-publication stages.

